# Mooney Face Image Processing in Deep Convolutional Neural Networks Compared to Humans

**DOI:** 10.1101/2022.03.21.485240

**Authors:** Astrid Zeman, Tim Leers, Hans Op de Beeck

## Abstract

Deep Convolutional Neural Networks (CNNs) are criticised for their reliance on local shape features and texture rather than global shape. We test whether CNNs are able to process global shape information in the absence of local shape cues and texture by testing their performance on Mooney stimuli, which are face images thresholded to binary values. More specifically, we assess whether CNNs classify these abstract stimuli as face-like, and whether they exhibit the face inversion effect (FIE), where upright stimuli are classified positively at a higher rate compared to inverted. We tested two standard networks, one (CaffeNet) trained for general object recognition and another trained specifically for facial recognition (DeepFace). We found that both networks perform perceptual completion and exhibit the FIE, which is present over all levels of specificity. By matching the false positive rate of CNNs to humans, we found that both networks performed closer to the human average (85.73% for upright, 57.25% for inverted) for both conditions (61.31% and 62.70% for upright, 48.61% and 42.26% for inverted, for CaffeNet and DeepFace respectively). Rank order correlation between CNNs and humans across individual stimuli shows a significant correlation in upright and inverted conditions, indicating a relationship in image difficulty between observers and the model. We conclude that in spite of the texture and local shape bias of CNNs, which makes their performance distinct from humans, they are still able to process object images holistically.

## Introduction

Humans are able to quickly recognise and categorise objects (Thorpe, Fize, & Marlot, 1996). The underlying neural system is conceptualised as an hierarchical information processing system that starts from a simple analysis of the visual input in terms of local features and ends with a high-level representation of complex object and category properties (Grill-Spector & Malach, 2004; DiCarlo, Zoccolan, & Rust, 2012). Deep convolutional neural networks (CNNs) have been developed with a similarly deep and hierarchical architecture, demonstrating success in various object recognition tasks. The representations developed in these networks through learning show interesting similarities with human brain and behaviour (Güçlü & Gerven, 2015; Khaligh-Razavi & Kriegeskorte, 2014; Kheradpisheh, Ghodrati, Ganjtabesh, & Masquelier, 2016; Yamins, et al., 2014). For example, CNNs represent visual object shape (Kubilius, Bracci, & Op de Beeck, 2016), independently from object category, which corresponds well with orthogonal representations of shape and category along the visual ventral stream (Zeman, Ritchie, Bracci, & Op de Beeck, 2020).

Despite their success in object recognition, deep convolutional neural networks (CNNs) reveal a shortcoming in processing global over local shape information (Baker, Lu, Erlikhman, & Kellman, 2018; Geirhos, et al., 2019; Brendel & Bethge, 2019; Xu, et al., 2018). Baker *et al.* (2018) found that the classification performance of CNNs was resilient to disruptions in global shape, but susceptible to local contour changes, an effect opposite to that of human observers. Brendel & Bethge (2019) argued that CNN performance is strongly correlated with local feature information by demonstrating that high classification performance in ImageNet (Russakovsky, et al., 2015) could be achieved by a bag-of-features network that had reduced patch size with disrupted spatial relationships between patches. Geirhos *et al.* (2019) assessed network classification using a cue conflict paradigm, where the texture of one object was superimposed over another object, and CNNs demonstrated a preference for object texture over shape, in contrast to humans. Xu *et al.* (2018) found that CNNs trained to identify faces are more sensitive to changes in texture compared to shape. These studies provide mounting evidence for an emphasis on local processing measures in CNNs, making their performance distinct from humans.

To interrogate the level of local versus global processing in CNNs, we assess their performance using a large battery of Mooney images, which are low-information images of faces that invoke face perception despite their high ambiguity (Mooney, 1957). Mooney images contain only isoluminant black and white areas and therefore have no texture and reduced local shape information. Global shape information processing is necessary for perceiving faces in these two-tone images, in which an observer must be able to integrate incomplete segments into a perceptual whole, known as “perceptual closure” (Wertheimer, 1923). The ability to separate Mooney faces from scrambled images is therefore a relevant index to assess this form of global shape processing. Recently, CNNs have been found to exhibit perceptual closure when trained on natural images (Kim, Reif, Wattenberg, & Bengio, 2019). CNNs that are randomly initialised or trained on random data do not exhibit closure, and closure ability is correlated with higher layers, peaking in the layer prior to prediction read-out (Kim, Reif, Wattenberg, & Bengio, 2019). Feed-forward networks display moderately high levels of pattern completion, which can be improved upon with the addition of recurrent connections to achieve human-level performance (Tang, et al., 2018). Given that CNNs are capable of closure and pattern completion, and are able to detect faces (Le, et al., 2012), it is logical that they are able to detect faces within Mooney images, which has indeed been established in a set of Mooney images artificially generated by a CNN (Ke, Yu, & Whitney, 2017).

Here we investigate closure ability in CNNs at a finer level than in previous studies, allowing us to establish a relationship with human perception of Mooney faces. One major aim is to test the effect of face inversion (Yin, 1969; Taubert, Apthorp, Aagten-Murphy, & Alais, 2011), which is one paradigm used to interrogate one form of global processing referred to as holistic processing (Piepers & Robbins, 2012), next to composite tasks (Young, Hellawell, & Hay, 1987) and part-whole tasks (Tanaka & Farah, 1993). Given that the parts are no longer accessible in Mooney faces, complicating the use of composite and part-whole tasks, the effect of inversion is the most obvious candidate paradigm to further test global processing in Mooney faces. Human observers show a clear face inversion effect (FIE) with Mooney faces, where a vertically mirrored Mooney image produces lower detection rates, slower reaction times and reduced activation in face-specific areas of the brain (Latinus & Taylor, 2005; Kanwisher, Tong, & Nakayama, 1998).

Another major aim is to investigate whether there is a correspondence between human and CNN performance in terms of which stimuli are more easily detected as containing a face. Previous studies in other domains have shown that patterns of errors can be similar between humans and CNNs, yet there is not always a tight relationship up to the image level (Lindsay, 2020; Rajalingham, et al., 2018). Here we make use of a recent extension of the original set of 50 Mooney faces (Mooney, 1957) to over 500 Mooney faces, with accompanying human observer accuracy and reaction times (Schwiedrzik, Melloni, & Schurger, 2018). With this larger dataset, we can thoroughly test CNNs and their ability to process local versus global shape information and compare directly to human behavioural ratings (which is absent from Ke, Yu, & Whitney, 2017).

In sum, we address the following questions. First, can deep networks detect faces in a large set of Mooney stimuli and demonstrate robustness in handling low information images that contain shadow, incomplete contours, no colour information, etc? Second, is there an inversion effect for Mooney faces in CNNs? Third, are the images that are difficult for humans also difficult for networks?

## Methods

### Stimuli

We use the Mooney face online dataset containing 96 scrambled, 504 upright and 504 inverted images from Schwiedrzik, Melloni, & Schurger (2018). Scrambled images contain contiguous regions with no sharp boundaries and were classified by humans to contain “no face” over 85% of the time in a pilot study (the false positive rate was higher during the actual study). Images were resized to 256 x 256 pixels prior to standard model input transformations. The online database of Schwiedrzik, Melloni, & Schurger (2018) has slightly fewer images than reported in their study, which were excluded due to copyright reasons. To ensure a fair comparison between human and CNN performance, we report only on data that relates directly to the images available online.

### CNNs

We test classification performance of two standard CNNs, one that has been trained for general object recognition (CaffeNet), and another that is trained specifically for facial recognition (DeepFace).

#### CaffeNet

– is an 8-layer, sequentially organised network with 5 convolutional layers followed by 3 fully-connected layers. CaffeNet is an implementation of AlexNet (Krizhevsky, Sutskever, & Hinton, 2012) in the Caffe Framework (Jia, et al., 2014). CaffeNet is trained for general object recognition using the ImageNet database (Russakovsky, et al., 2015) and to distinguish between 1000 classes that do not explicitly contain faces.

We assess positive performance of CaffeNet by taking the average of its classification of images into 2 categories that may be considered as face-like, namely Mask and Ski Mask. These face-like categories are most likely to contain a feature configuration with two-eyes above a mouth, and sometimes contain human faces within the actual images (wearing the mask). For classification performance, we take the top 5 model predictions, where chance level is calculated to be 1.002% using the following reasoning. Chance level for selecting one of the 2 face-like categories as the Top 1 selection is simply 2/1000 or 0.2%. To calculate chance level for the Top 2 results, we add 0.2% with the possibility of 2 out of 999 remaining selections (having non-replacement of categories), assuming that one of the 2 possible categories was not selected as the top 1 result. We repeat this process to calculate the Top 5 chance level, which is 2/1000 + 2/999 + 2/998 + 2/997 + 2/996 = 1.002%.

#### DeepFace

– is a serial network with 6 parameterised layers (4 convolutional followed by 2 fully-connected) and 3 max pooling layers that follow every convolutional layer (Chen, 2016). DeepFace performs face detection, alignment and recognition, and is trained on the CASIA-WebFace database. We assess positive performance of DeepFace using binary recognition, where chance level is 50%.

### ROC curve and rank-order correlations

A Receiver Operating Characteristic or ROC curve is created by plotting the true positive rate (sensitivity) against the false positive rate at different threshold levels. For our purposes, an ROC curve allows us to determine how well each CNN is able to detect faces in the Mooney images under differing levels of specificity. To construct the ROC curves, we take each image and run a forward pass through the model and read out the probability predicted by the model that it belongs to the face class. This probability is thresholded so that the images associated with a probability above the threshold are classified as faces and the other images as nonfaces. Then, under different threshold levels, we determine the percentage of face images that are classified as belonging to the face class as well as the percentage of nonface images that are classified as a face. To calculate the rank-order correlations for each model, we also use the probability scores that are generated for each image by each model. For CaffeNet, we take the average of the two probabilities that the image belongs to two of the face-like classes, and use this to obtain the rank order.

## Results

### Perceptual Completion and the FIE in CNNs

We first tested the networks (CaffeNet, DeepFace) in their ability to classify an image as containing a face (in the case of DeepFace) or a face-like object (in the case of CaffeNet), for each of the three conditions (upright, inverted and scrambled). CaffeNet was evaluated using the top 5 category ranking, for two face-like categories (Mask or Ski-mask). DeepFace was evaluated based on the binary classification of whether each image contained a face or not. Average human accuracy was 85.73% for upright Mooney faces, and 57.25% for inverted, calculated from raw data in Schwiedrzik, Melloni, & Schurger (2018). On average, humans recognised faces 27.36% of the time in the 96 scrambled images, determining the average false positive level.

If the networks are able to perform perceptual completion of the Mooney face stimuli, we expect to see higher classification accuracy for upright versus scrambled images. If the networks exhibit an FIE, we expect to see higher performance for upright versus inverted images. Figure 1 illustrates the classification results for the two CNNs. Both networks were able to perform perceptual completion, showing upright performance as higher than scrambled (CaffeNet: difference of 28.03%, DeepFace: 28.92). Both CNNs exhibited an FIE, with upright performance being higher than inverted (CaffeNet: 15.48% difference between upright and inverted, DeepFace: 23.41 difference). Both networks showed results qualitatively consistent with human performance – with both upright and inverted accuracy levels being above the false positive (scrambled) rate. Even though both CNNs showed higher performance for upright (Caffenet: 47.82%, DeepFace: 38.29%) versus inverted (CaffeNet: 32.34%, DeepFace: 14.88%) images, this was still below the average human response of 85.73% and 57.25% for these same images, however the false positive rate was also lower for the CNNs (CaffeNet: 19.79%, DeepFace: 9.38%) compared to humans (27.36%).

**Figure 1:**
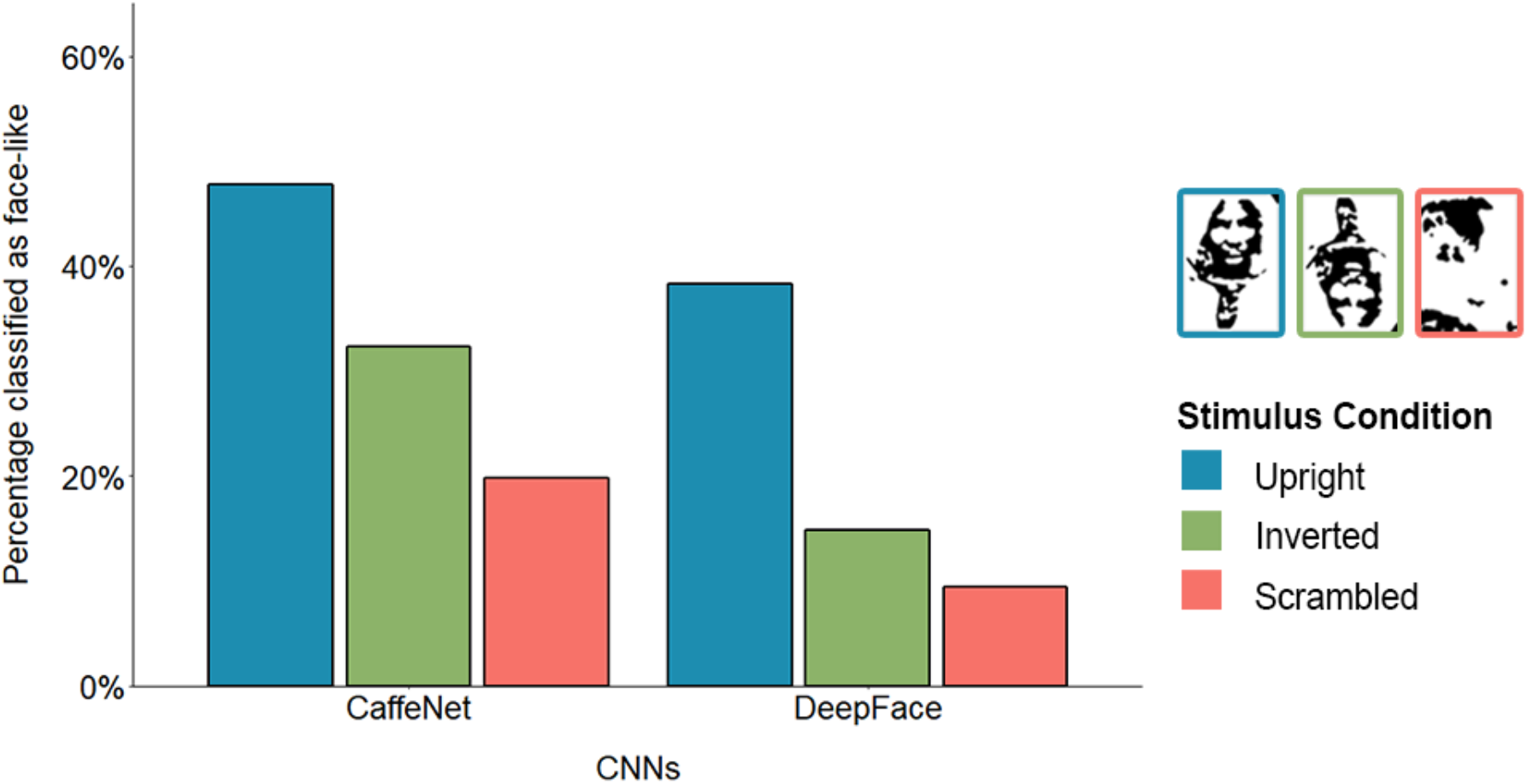
Classification in CNNs of images as faces or as face-like, in object-trained CaffeNet (left) and face-trained DeepFace (right)

Considering that the false positive rate (scrambled image performance) for both CNNs was below the human average, we plotted an ROC curve to better determine the upright versus inverted performance of each CNN when the false positive rate matched humans (see Methods). CaffeNet ROC curves for upright (blue) and inverted (green) conditions are shown in Figure 2, against human performance taken from Schwiedrzik, Melloni, & Schurger (2018). Average human accuracy levels are indicated by asterisks at the average false positive mark, and individual subject performance is indicated by squares. To plot CaffeNet performance, we measured the model’s predicted probability scores generated for the two face-like categories (Mask and Ski Mask) for each stimulus condition and averaged these. We then adjusted the detection threshold to determine the true positive rate (upright or inverted) versus the false positive rate (scrambled). When the false positive rate of CaffeNet was 27.08% (roughly equivalent to the human false positive rate of 27.36%), upright performance was 61.31% and inverted was 48.61. These values are now closer to the average human performance levels of 85.73% for upright and 57.25% for inverted (illustrated by the asterisks in Figure 2), although they are still not as high as human levels. For DeepFace, when the false positive rate was 27.08%, upright performance was 62.70% and inverted was 42.26%. One interesting feature to note is that for both networks, upright performance is always higher than inverted, demonstrating an FIE over all false positive levels. The size of the inversion effect remains roughly consistent across different levels of specificity.

**Figure 2:**
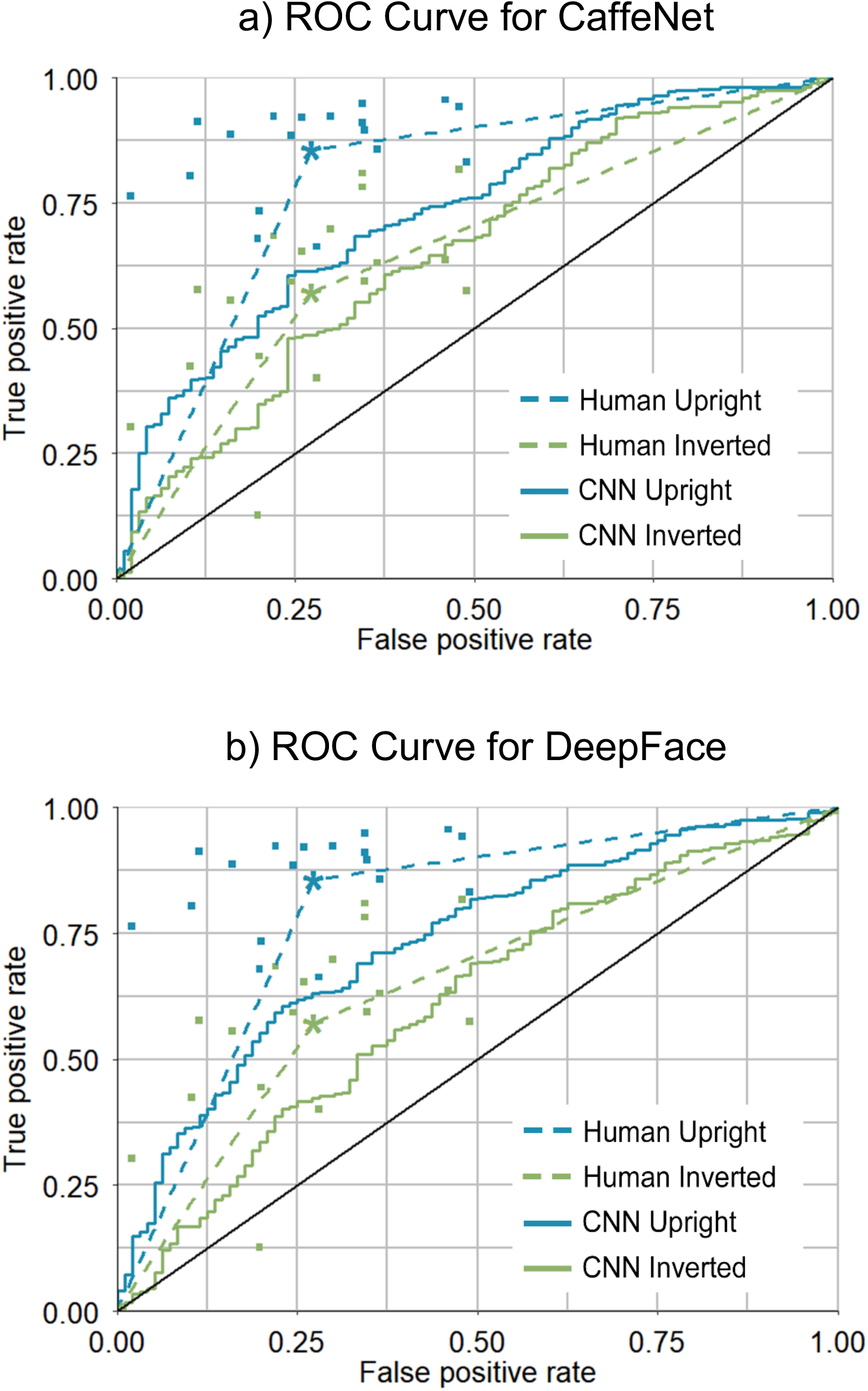
ROC curves for a) CaffeNet (top) and b) DeepFace (bottom) (solid lines), plotted against human performance (dashed lines) for upright (blue) versus inverted (green) images. Asterisks represent average human performance, and squares represent individual subjects. The black diagonal represents chance level.

### Rank order correlation between CNNs and humans

We tested the rank order correlation of CNN accuracy with human accuracy levels for upright versus inverted conditions. CaffeNet accuracy was calculated using the average prediction probabilities of Mask and Ski Mask categories. The noise ceiling was calculated from human performance using split-half cross validation across two subgroups of human participants. Results are shown in Table 1. Before Noise Correction (NC), Spearman correlation for CaffeNet was *ρ* = 0.26685 (*ρ* < 0.0001) for upright images and *ρ* = 0.17241 (*ρ* = 0.0001) for inverted. After NC, upright correlation was *ρ* = 0.31727 and inverted was *ρ* = 0.20814. For DeepFace, before NC, Spearman correlation was *ρ* = 0.28807 (*ρ* < 0.0001) for upright images and *ρ* = 0.21943 (*ρ* < 0.0001) for inverted. After NC, Spearman correlation was *ρ* = 0.34250 (*ρ* < 0.0001) for upright images and *ρ* = 0.26491 (*ρ* = 0.014) for inverted. These values demonstrate a significant, although not a large relationship, between image difficulty for humans and CNNs. The strength of rank order correlation appears to also correlate with performance, given that higher rho values correspond with higher accuracy levels, in agreement with Yamins, *et al.* (2014).

**Table 1:** Rank order correlation between CNNs and human accuracy. NC is Noise Corrected.

|  | Spearman's<br>rho | NC Spearman's<br>rho | NC 95%<br>Low CI | NC 95%<br>High CI | Noise Ceiling |
| --- | --- | --- | --- | --- | --- |
| CaffeNet<br>Upright | 0.26685 | 0.31727 | 0.21589 | 0.41375 | 0.84108 |
| CaffeNet<br>Inverted | 0.17241 | 0.20814 | 0.10479 | 0.30964 | 0.82832 |
| DeepFace<br>Upright | 0.28807 | 0.34250 | 0.24403 | 0.43991 | 0.84108 |
| DeepFace<br>Inverted | 0.21943 | 0.26491 | 0.16121 | 0.36135 | 0.82832 |

## Discussion & Conclusion

In this paper we examined two standard CNN models and their ability to detect faces in ambiguous Mooney images. We first analysed network performance using the default discrimination levels for each CNN and found that the false positive rate for each model (using scrambled images) was lower than in human observers. We then examined the face detection levels of each network under differing levels of specificity, to determine levels of performance when the false positive rate was matched to humans. Finally, with each model, we ranked images in their predicted probability to contain a face and correlated this with human observers to quantify any overlapping similarity in image difficulty. From these experiments, we present three main findings: 1. Deep networks are able to detect faces in Mooney stimuli, demonstrating robust detection in these low information images; 2. CNNs exhibit a face inversion effect with Mooney faces, which is evident across all degrees of specificity; and 3. Mooney images that are difficult for humans are also difficult for CNNs, as demonstrated by a significant rank order correlation between CNNs and humans for upright and inverted images.

In this study, we selected two CNNs that were trained for very different purposes using very different datasets. One network (CaffeNet) was trained for general object recognition, whereas the other (DeepFace) was trained for face recognition, which is more relevant to this task. Despite the large differences in training datasets, both networks, surprisingly, behaved quite similarly. The large agreement between both networks is most evident in the ROC curves, which show similar levels of performance. Note that neither network was trained using Mooney images, which we would expect would improve performance and also potentially improve the correlation with human observers. The results that we present here showcase the generalisability and robustness of these networks.

While we do not claim that these networks are equivalent to human object and face processing, it is clear that they are capable of similar feats, albeit to a lesser extent. For example, while CNNs exhibit a bias for texture over shape (Baker, Lu, Erlikhman, & Kellman, 2018; Geirhos, et al., 2019; Brendel & Bethge, 2019; Xu, et al., 2018), and a bias for local over global shape (Baker, Lu, Erlikhman, & Kellman, 2018;), they are still able to classify objects with no texture information, in the form of silhouettes (Kubilius, Bracci, & Op de Beeck, 2016). Nevertheless, when controlling for local shape information, as tested by the GIST model, CNNs represent global shape information in multiple network layers (Zeman, Ritchie, Bracci, & Op de Beeck, 2020). We hypothesize that these models could achieve better levels of performance, on par with humans, with the integration of recurrent connections at higher network layers, as implemented in Tang et al. (2018). In addition, we also expect effects of changes in training history that go beyond the differences in training regime between the two networks that we have tested here. Shape or texture preferences in networks can be adjusted via the training regime of networks, as demonstrated recently by Geirhos *et al.* (2019). The authors conclude that texture bias can be reduced, or even overcome, with the adjustment of weights within networks, to better promote global processing. In relation to our work, we can see that although there is a difference between human and CNN performance, the difference is in weighting schemes, rather than representational capabilities. To reduce the gap between human and CNN performance, we also suggest adopting a similar training regime as described by Geirhos *et al.* (2019).

We conducted ROC analyses to determine the discrimination levels of each network under varying degrees of false positive rates, particularly noting when the level was matched to that of humans. At all false positive levels, we found upright discrimination was higher than inverted, demonstrating the robustness of the inversion effect. Effects in CNNs were in the same direction as humans, but the effect sizes were smaller in CNNs. When examining CNN versus human performance more closely using rank order correlations, the correlations reached a certain level but were definitely not at ceiling. This suggests that there are still differences between CNNs and humans at the image level.

Here we show that CNNs can classify low information images of quite complex visual stimuli to perform perceptual completion on images with no texture and no obvious local cues to dissociate face from nonface images. CNNs also exhibit holistic processing on these zero texture images, at all layers of specificity, as demonstrated by the inversion effect. In conclusion, despite the preference of CNNs to take into account texture over shape, which differs from humans, they still process object images in a holistic manner, demonstrating both perceptual completion and the inversion effect.

## Acknowledgements

We thank Caspar Schwiedrzik for providing additional raw data and help in answering questions regarding his study. This work was funded by grant C14/16/031 of the KU Leuven Research Council.

## Notes

### Competing Interest Statement

The authors have declared no competing interest.

